# Ketolysis is Required for the Proper Development and Function of the Somatosensory Nervous System

**DOI:** 10.1101/2023.01.11.523492

**Authors:** Jonathan Enders, Jarrid Jack, Sarah Thomas, Paige Lynch, Sarah Lasnier, Xin Cao, M Taylor Swanson, Janelle M Ryals, John P. Thyfault, Patrycja Puchalska, Peter A. Crawford, Douglas E Wright

## Abstract

Ketogenic diets are emerging as protective interventions in preclinical and clinical models of somatosensory nervous system disorders. Additionally, dysregulation of succinyl-CoA 3-oxoacid CoA-transferase 1 (SCOT, encoded by *Oxct1*), the fate-committing enzyme in mitochondrial ketolysis, has recently been described in Friedreich’s ataxia and amyotrophic lateral sclerosis. However, the contribution of ketone metabolism in the normal development and function of the somatosensory nervous system remains poorly characterized. We generated sensory neuron-specific, Advillin-Cre knockout of SCOT (Adv-KO-SCOT) mice and characterized the structure and function of their somatosensory system. We used histological techniques to assess sensory neuronal populations, myelination, and skin and spinal dorsal horn innervation. We also examined cutaneous and proprioceptive sensory behaviors with the von Frey test, radiant heat assay, rotarod, and grid-walk tests. Adv-KO-SCOT mice exhibited myelination deficits, altered morphology of putative Aδ soma from the dorsal root ganglion, reduced cutaneous innervation, and abnormal innervation of the spinal dorsal horn compared to wildtype mice. Synapsin 1-Cre-driven knockout of *Oxct1* confirmed deficits in epidermal innervation following a loss of ketone oxidation. Loss of peripheral axonal ketolysis was further associated with proprioceptive deficits, yet Adv-KO-SCOT mice did not exhibit drastically altered cutaneous mechanical and thermal thresholds. Knockout of *Oxct1* in peripheral sensory neurons resulted in histological abnormalities and severe proprioceptive deficits in mice. We conclude that ketone metabolism is essential for the development of the somatosensory nervous system. These findings also suggest that decreased ketone oxidation in the somatosensory nervous system may explain the neurological symptoms of Friedreich’s ataxia.

## Introduction

Ketolysis—the catabolism of β-hydroxybutyrate (β-HB) and acetoacetate for fuel—is known to contribute to the development and health of the nervous system. Ketone utilization is highest during neonatal periods, with β-HB uptake and activity of β-HB dehydrogenase 1 (BDH1) decreasing throughout development and into adulthood (Kraus et al., 1974, Bilger and Nehlig, 1992, Nehlig et al., 1991). Acetoacetate prevents hippocampal lesions caused by the inhibition of glycolysis in Wistar rats and increased adenosine triphosphate (ATP) production (Massieu et al., 2003). Likewise, β-HB is neuroprotective and supports synaptic function following glycolytic inhibition in hippocampal slices from rats postnatal day 30 and earlier (Izumi et al., 1998). In disease states, ketogenic diets are neuroprotective in pediatric epilepsy (Sourbron et al., 2020, Henderson et al., 2006), and their use has led to improvements in Alzheimer’s disease, cognitive impairment (Taylor et al., 2018), and migraine (Bongiovanni et al., 2021, Di Lorenzo et al., 2019). Together, these findings highlight the importance of metabolic plasticity between glycolysis and ketolysis in the developing and diseased central nervous system.

Little is known about ketone metabolism’s contribution to somatosensory nervous system health. Ketone bodies are brought into peripheral tissues by the monocarboxylate transporters (Hugo et al., 2012), after which they are utilized for metabolic and signaling functions (Puchalska and Crawford, 2017, Puchalska and Crawford, 2021). The obligate reaction during ketolysis is the addition of coenzyme A to acetoacetate catalyzed by succinyl-CoA 3-oxoacid CoA-transferase 1 (SCOT, encoded by *Oxct1*). *Oxct1* mRNA and SCOT protein expression are downregulated in mouse models of neurological diseases, including amyotrophic lateral sclerosis (Szelechowski et al., 2018) and Friedreich’s ataxia (Dong et al., 2022). Ketone incorporation in lipid synthesis is disrupted in the sciatic nerve of Trembler mouse models of Charcot-Marie-Tooth (Clouet and Bourre, 1988). We previously published that ketone bodies support neurite outgrowth from dissociated dorsal root ganglia (Cooper et al., 2018b). We also recently showed that ketone bodies are further protective in painful neuropathies by contributing to the detoxification of reactive dicarbonyls, such as methylglyoxal, that are elevated in diabetic neuropathy and directly cause neuronal dysfunction and pain (Enders et al., 2022b). Likewise, our works showed that a ketogenic diet supported the regeneration of lost intraepidermal nerve fibers in a model of peripheral diabetic neuropathy (Enders et al., 2022a) and reversed pain-like behaviors in mouse models of type 1 diabetes (Enders et al., 2022a) and prediabetes (Cooper et al., 2018b). Additionally, sciatic nerve mitochondria from mice fed a ketogenic diet produce less reactive oxygen species (Cooper et al., 2018a). Together, these prior findings suggest ketone metabolism likely plays a vital role in the protective effects of a ketogenic diet in the peripheral nerve.

To understand the contributions of ketone metabolism in the somatosensory nervous system, we used a Cre-lox system to generate a peripheral sensory neuron-specific knockout of *Oxct1,* referred to as sensory neuron, Advillin-Cre knockout of SCOT (Adv-KO-SCOT) mice. We tested the hypothesis that ketone body metabolism or signaling was critical for somatosensory nervous system development by exploring potential changes in sensory neuronal phenotypes, peripheral axon integrity, and somatosensation in Adv-KO-SCOT mice. Our results suggest that ketone metabolism in sensory neurons is required for normal epidermal innervation and myelination of large peripheral axons. Sensory behaviors largely remain unchanged in Adv-KO-SCOT mice; however, these animals’ response to intraplantar capsaicin was diminished and they demonstrated proprioceptive deficits. These findings provide important insight into processes underlying ketone metabolism in the peripheral nerves. Moreover, these results provide an essential framework for understanding the potential therapeutic effects of a ketogenic diet on somatosensation and pain.

## Materials and Methods

### Animals and Breeding

All animal work was performed following review and approval by the Institutional Animal Care and Use Committee of Kansas University Medical Center. All mice were maintained on a 12:12 light: dark cycle in the Kansas University Medical Center animal research facility. In studies containing a ketogenic diet, mice were given *ad libitum* access to a ketogenic diet (TD.96355; Envigo, 90.5% fat, 9.2% protein, and 0.3% carbohydrate by kcal), which was replaced twice weekly. C57Bl6 mice carrying the *Oxct1^flox/flox^*allele and Synapsin-1 (*Syn1*)*^+/Cre^Oxct1^flox/flox^* mice were generated previously (Cotter et al., 2013). *Oxct1^flox/flox^*mice were crossed with *Advillin^Cre/+^*or *Syn1^+/Cre^*C57Bl6 mice (Jackson Laboratory, Bar Harbor, ME) to generate *Oxct1^flox/+^Advillin^Cre/+^*and *Oxct1^flox/+^Syn1^+/Cre^*mice. These mice were bred to *Oxct1^flox/flox^*mice, generating *Oxct1^flox/flox^Advillin^Cre/+^* and *Oxct1^flox/flox^Syn1^+/Cre^* mice. Male *Oxct1^flox/flox^Advillin^Cre/+^* and *Oxct1^flox/flox^Syn1^+/Cre^* mice were bred to *Oxct1^flox/flox^*females to produce sensory neuron, Advillin-Cre knockout of SCOT (Adv-KO-SCOT) mice and Syn1-Cre mice, respectively. Sensory neuronal knockout of *Oxct1* was confirmed by immunofluorescent microscopy of the dorsal root ganglion (DRG) (**Figure 1**). We primarily used Adv-KO-SCOT mice throughout this work and included Syn1-Cre mice as evidence that the observed effects were due to Oxct1 knockout and unrelated to Cre toxicity.

**Figure 1.**
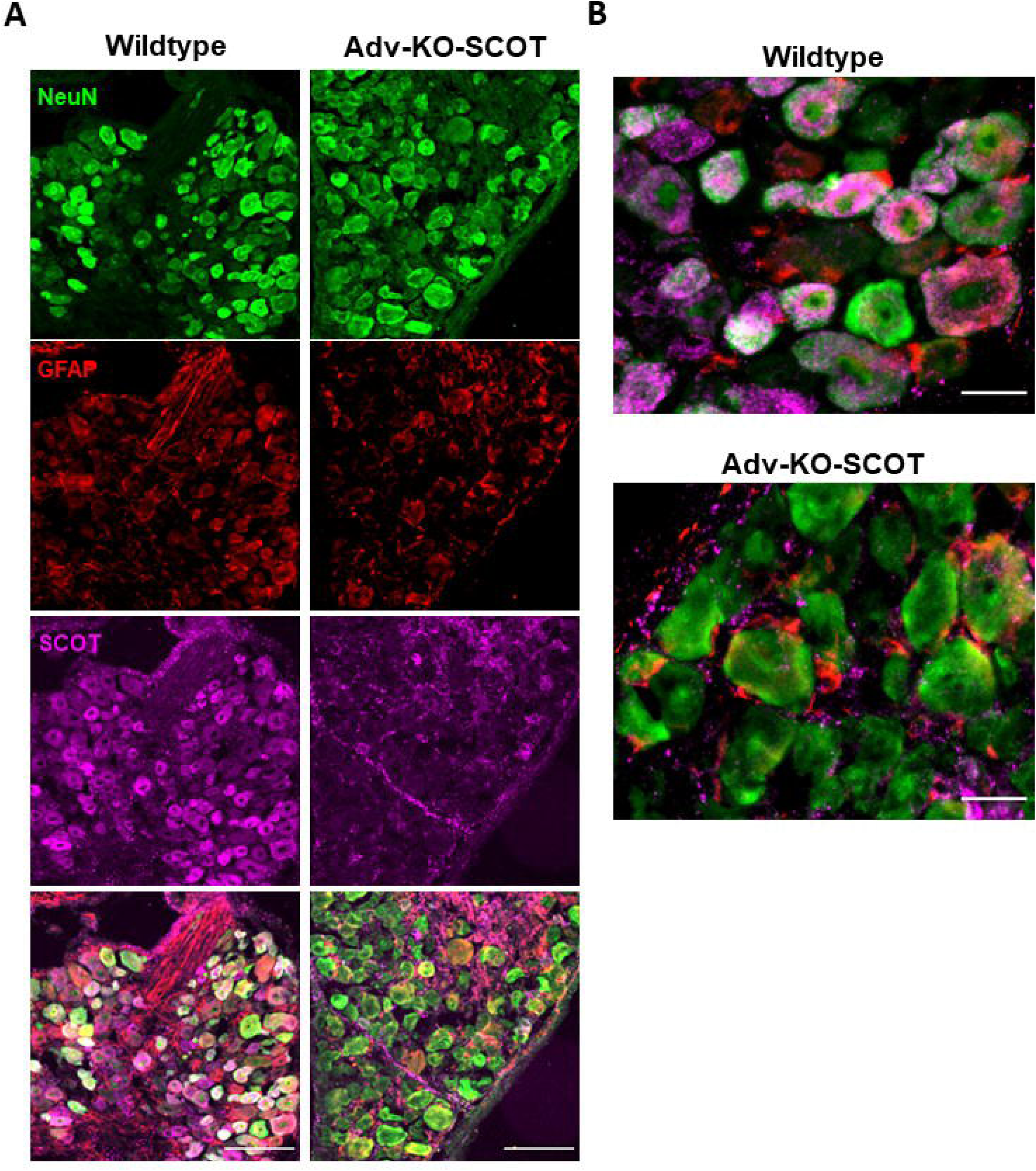
Adv-KO-SCOT mice lack expression of *Oxct1* in the dorsal root ganglia. (A) Representative micrographs of dorsal root ganglia (DRG) stained from wildtype (WT) and Sensory nerve, Advillin-Cre Knockout of SCOT (Adv-KO-SCOT) mice, taken at 10X. While most NeuN^+^ cells in wildtype DRG are also SCOT^+^, virtually no neurons in Adv-KO-SCOT mice contain SCOT. Scale bar represents 100 μm. (B) Representative micrographs of wildtype and Adv-KO-SCOT DRG with the same stain taken at 40X. Virtually no GFAP^+^ cells stained positively for SCOT in either genotype. Scale bar represents 25 μm.

### Blood Measurements

Fasting blood glucose was determined as described by Groover et al. (Groover et al., 2013). Briefly, mice were fasted for 3 hours before drawing 20 μL blood from the tail vein by clipping off the tip of the tail or the resulting scab in subsequent blood draws. We used Vaseline during the blood draw to avoid skin abrasion and improve scab healing. All standards and samples were mixed with molecular H_2_O, ZnSO_4_ (Sigma), and Ba(OH)_2_ (Sigma) and centrifuged at 13000 rpm and 4 °C for 5 minutes. The supernatant was incubated for 30 minutes at 37 °C with color reagent (PGO capsule, Sigma; o-Dianisidine Dihydrochloride, Sigma; in molecular water). A plate reader (SpectraMax M5) determined the absorbance of standards and samples at 450 nm. Blood ketones were measured using a hand-held blood monitor and β-hydroxybutyrate blood ketone strips (β-Ketone blood test strips, Abbott Laboratories, Chicago, IL; Precision Xtra, Abbott Laboratories) after a 3-hour fast.

### Cutaneous Sensory Behavior

Thresholds to mechanical and thermal stimulation were collected every 4 weeks in Adv-KO-SCOT mice starting at six weeks of age using Von Frey microfilaments and a Hargreaves Analgesiometer, respectively. Before data collection, mice were acclimated to testing areas for 30 minutes and the mesh table or thermal testing apparatus for 30 minutes on two occasions, separated by 24 hours. Finally, mice were acclimated to the testing area, mesh table, and thermal testing apparatus for 30 minutes each before collecting data.

Different Von Frey monofilaments were applied for one second to the plantar surface of the hind paw following the “up-down” method (Chaplan et al., 1994). Mice were observed for either a negative or a positive response, and the mechanical withdrawal threshold was calculated after five consecutive responses following the first positive response. Thermal sensitivity was determined by radiant heat assay using a thermal testing apparatus. A 4.2 V radiant heat source was applied to the plantar surface of the hind paw, and latency to withdrawal was recorded three times.

Mice were acclimated to a clear plastic cage without bedding for 5 minutes before capsaicin injection. Following intraplantar injection, mice were replaced in the cage. Capsaicin was diluted in sterile saline with 0.5% Tween 20 to a working concentration of 0.1 μg/μL and delivered by 20 μL subcutaneous injection to the right hind paw (2 μg). A blinded investigator then observed the mouse for 5 minutes following injection and recorded the number of nocifensive events the animal displayed (licking, lifting, biting, guarding, etc.) and the total time spent engaged in nocifensive behavior.

### Proprioceptive Sensory Behavior

Proprioception was measured at six weeks of age by a Rotorod test. Mice were acclimated to the rotarod via training sessions the day before data collection. On the day of data collection, mice were placed on the Rotarod treadmill (Med Associates, St Albans, VT). The rod of 32mm diameter, elevated 16 cm above the lab bench, was set to ramp from 4 rpm to 40 rpm over a 300-sec duration as the mouse locomoted on the rod. Three trials were conducted with an intertrial interval of not less than 5 minutes, assessing an average latency to fall.

For the grid-walk test, mice were placed on a 35cm x 35cm wire mesh elevated 45cm above a lab bench with 1 cm^2^ openings. During testing, mice were allowed to walk freely on the grid for 5 minutes while being filmed from grid level using a portable camera. Following testing, videos were reviewed by a blinded observer and analyzed for time spent walking, forepaw and hind paw slips, and total paw slips. Paw slips were defined as an extension of the limb through a wire opening to at least the ankle joint. Paw slips were normalized to time walking (slips/minute).

### Tissue Processing and Immunofluorescent Microscopy

Mice were sacrificed by inhalation of isoflurane and decapitation. Dorsal root ganglia (DRG), footpads, and sciatic nerve were post-fixed overnight in 4% paraformaldehyde, Zamboni’s fixative, and 2.5% glutaraldehyde with 2% paraformaldehyde, respectively. Fixed tissues were then rinsed with PBS. DRG, spinal cord, and footpad were soaked in 30% sucrose, rinsed, and then frozen in Optimal Cutting Temperature Compound (Sekura Tissue-Tek). Gastrocnemius were dissected away from the right hindlimb. The soleus was removed, and gastrocnemius were stretched and fresh-frozen on a penny.

### Dorsal Root Ganglia SCOT Detection and Sensory Neuronal Population Assessment

Eight-micron sections of L4-6 DRG were blocked for two hours in Superblock (ThermoFisher; Grand Island, NY), 1.5% Normal Donkey Serum, 0.5% Porcine Gelatin, and 0.5% Triton X-100 (Sigma) at room temperature. For SCOT detection, slides were incubated overnight with goat anti-neuronal nuclear protein (NeuN) (1:1000, Novus Biologicals; Centennial, CO), chicken anti-glial fibrillary associated protein (GFAP) (1:1000, Abcam; Boston, MA), and rabbit anti-SCOT (1:250, Abcam; Boston, MA). The next morning, slides were washed twice with PBS and stained with AlexaFluor 488-tagged donkey-anti-goat (Molecular Probes, 1:100), Cy3-tagged donkey-anti-chicken (Jackson Immuno, 1:200), and AlexaFluor 647-tagged donkey-anti-rabbit (Molecular Probes, 1:1000) secondary antibodies for one hour. Slides were then washed twice with PBS and coverslipped. Fluorescent images were taken at 20x with a Nikon Eclipse Ti2 inverted microscope.

Slides were incubated overnight with anti-TrkA (goat, R&D Technologies, 1:250) and anti-NF-H (rabbit, EMD Millipore, 1:1000) antibodies to assess various sensory neuronal populations. Slides were washed twice with phosphate-buffered saline for 5 minutes and incubated with Alexa Fluor 555-tagged donkey-anti-goat (Molecular Probes, 1:1000) and Alexa Fluor 647-tagged donkey-anti-rabbit (Molecular Probes, 1:1000) secondary antibodies for 1 hour. Slides were rinsed twice with PBS and incubated in freshly prepared Alexa Fluor 488-tagged IB4 (Invitrogen, 10μg/mL in 1mM CaCl_2_) for 10 minutes. Slides were rinsed twice in PBS and coverslipped before imaging with a Nikon Eclipse 90i microscope at 20x. At least five images were taken from each DRG. Neuron soma area and Feret diameter were determined by measurements with ImageJ software, and neurons were counted manually.

### Intraepidermal Nerve Fiber Quantification

Thirty-micron sections of the footpad were blocked for two hours in Superblock (ThermoFisher; Grand Island, NY), 1.5% Normal Donkey Serum, 0.5% Porcine Gelatin, and 0.5% Triton X-100 (Sigma) at room temperature. Slides were incubated overnight with rabbit anti-post gene product 9.5 (PGP9.5) (1:1500, UCHL1; ProteinTech; Rosemont, IL) and goat α-TrkA (1:250; R&D Systems, Minneapolis, MN). Slides were incubated with Alexa Fluor 555 tagged donkey-α-rabbit secondary antibody (1:1000; Molecular Probes) and Alexa Fluor 488-tagged donkey-α-goat secondary antibody (1:1000; Molecular Probes) for one hour, then imaged with a Nikon Eclipse 90i microscope using a 20X objective. ImageJ software was used to measure the length of the dermal-epidermal junction. IENF was quantified as the number of fibers crossing that junction and expressed as the number of fibers per millimeter. Nine images were taken per slide, and the average IENF density for each mouse was used for statistical analyses.

### Quantification of Myelination

Sciatic nerves were dissected out of 6-week-old WT and Adv-KO-SCOT mice, post-fixed in 2.5% glutaraldehyde and 2% paraformaldehyde overnight at 4 °C and rinsed with phosphate-buffered saline. Tissue was then embedded in paraffin blocks, semithin (1 μm) cross sections were cut by microtome, and sections were stained with toluidine blue before coverslipping. Brightfield images were taken at 30x with a Nikon Eclipse Ti2 inverted microscope. The “GRatio” plugin was used with ImageJ to randomly select and measure the area and perimeter of axons and their associated myelin from each image. Thin sections (70-85 μm) were cut with a DiATOME diamond knife, post-stained with uranyl acetate and lead citrate, and imaged with a JEOL JEM-1400 transmission electron microscope (TEM) at 100 kV. At least five images were taken of each sample at 1000x and 2000x. Dysmyelination was defined as invagination, separation from the axoplasm, and unraveling of the myelin (**Figure 4E-F**). These events were quantified as a frequency of dysmyelinated axons among all myelinated axons completely captured in a field of view by a blinded observer. Remak bundle occupancy was quantified by counting the total number of unmyelinated fibers within Remak bundles completely captured in a field of view normalized to the number of Remak bundles within that view.

### Quantification of Spinal Dorsal Horn Innervation

The lumbar enlargement of the spinal cord was dissected out of 6-week-old WT and Adv-KO-SCOT mice, post-fixed in 4% paraformaldehyde and soaked in 30% sucrose. 30 μm sections were blocked for two hours in Superblock, 1.5% Normal Donkey Serum, 0.5% Porcine Gelatin, and 0.5% Triton X-100 at room temperature. Sections were stained overnight with goat anti-TrkA antibody (1:250). Sections were washed with phosphate-buffered saline, stained with freshly prepared Alexa Fluor 488-tagged IB4 (10μg/mL in 1mM CaCl_2_) for 10 minutes, washed again, and stained with Alexa Fluor 555-tagged donkey-α-goat secondary antibody (Molecular Probes, 1:1000) for 1 hour. Sections were coverslipped and imaged with a Nikon Eclipse Ti2 inverted microscope at 10x. The area of TrkA-positive innervation of lamina I and IB4-positive innervation of lamina IIo were quantified by ImageJ. A blinded observer manually counted the number of TrkA-positive projections extending through lamina II into lamina III or deeper. At least five images of the dorsal horn of the spinal cord were taken for each mouse, and the average TrkA^+^ and IB4^+^ areas and the number of deeper TrkA^+^-projections were used for statistical analysis.

### Assessment of Muscle Spindles

Serial 90 μm sections were cut from fresh-frozen gastrocnemius dissected from six-week-old WT and Adv-KO-SCOT mice. Sections were blocked overnight in Superblock, 1.5% Normal Donkey Serum, 0.5% Porcine Gelatin, and 0.5% Triton X-100 at room temperature. Sections were incubated with mouse anti-myosin heavy chain, slow developmental (S46, 1:50; Developmental Studies Hybridoma Bank; Iowa City, IA) and rabbit anti-NF-H (1:500; EMD Millipore; Burlington, MA) for 24 hours at room temperature. Slides were washed twice with filtered PBS and then incubated with AlexaFluor 555-tagged donkey-anti-rabbit (Molecular Probes, 1:1000) and AlexaFluor 488-tagged donkey-anti-mouse (Molecular Probes, 1:1000) secondary antibodies and Hoescht 33342 stain (Molecular Probes, 1:2000) for three hours at room temperature. Slides were washed again twice with filtered PBS. Slides were coverslipped and imaged with a Nikon Eclipse Ti2 inverted microscope at 40x. Two to seven muscle spindles were imaged per animal. For each spindle, the average width of three or more axonal rotations and the average distance between axonal rotations were measured by ImageJ.

### Statistical Analyses

All statistical analyses were performed using R version 3.6.2 and packages “Rmisc”, “car”, and “ggpubr”. All analyses for which data were collected over time were performed using a mixed-model analysis of variance (ANOVA) with repeated measures. All other analyses were performed using a two-way ANOVA or Student’s t-test, as appropriate. Data were analyzed posthoc by pairwise t-test or Tukey’s Honest Significant Difference (HSD) as indicated. Data are presented as mean +/- standard error of the mean in bar and line charts or as median with interquartile range in violin plots. Type I error rate was set at 0.05 except where noted due to a Bonferroni correction for multiple comparisons.

## Results

### Adv-KO-SCOT mice have nearly complete ablation of SCOT in dorsal root ganglia neurons

We confirmed the loss of *Oxct1* in sensory neurons using immunofluorescent microscopy. Neuronal soma (marked by NeuN) in the dorsal root ganglia (DRG) were stained positively for SCOT in *Oxct^flox/flox^Advillin^+/+^*mice (wildtype mice, hereafter) (**Figure 1**). Adv-KO-SCOT mice were negative for SCOT staining in the neuronal soma of the DRG, indicating the absence of the enzyme from peripheral sensory neurons. Notably, GFAP^+^ cells were also devoid of SCOT staining in wildtype and Adv-KO-SCOT DRG.

### Sensory neuronal knockout of Oxct1 has minimal effect on whole-body metabolism

Neuronal knockout of *Oxct1* has been associated with the dysregulation of blood glucose and serum ketones (Cotter et al., 2013). Given the effect of metabolic syndrome and diabetes on peripheral nervous system health, we characterized metabolic markers in wildtype and Adv-KO-SCOT mice. Adv-KO-SCOT mice were modestly but significantly heavier than wildtype mice (**Figure 2A**; N-way mixed-model ANOVA with repeated measures, genotype: p = 0.00176). We detected a sex difference in the effect of genotype (genotype-sex interaction: p = 7.28e^-5^). Adv-KO-SCOT males trended toward reduced body weight compared to their wildtype littermates (**Figure 2B**; Tukey’s *post hoc*, p = 0.055), while Adv-KO-SCOT females were significantly heavier than their wildtype littermates (**Figure 2C**; Tukey’s *post hoc*, p = 0.0041). We did not detect sex differences in fasting blood glucose or circulating ketones and thus pooled and reanalyzed the data. There was no significant difference in fasting blood glucose between chow-fed wildtype and Adv-KO-SCOT mice (**Figure 2D**; N-way mixed-model ANOVA with repeated measures, genotype: p = 0.103, genotype-time interaction: p = 0.508), nor were there differences in the abundance of circulating β-HB (**Figure 2E**; N-way mixed-model ANOVA with repeated measures, genotype: p = 0.506, genotype-time interaction: p = 0.701).

**Figure 2.**
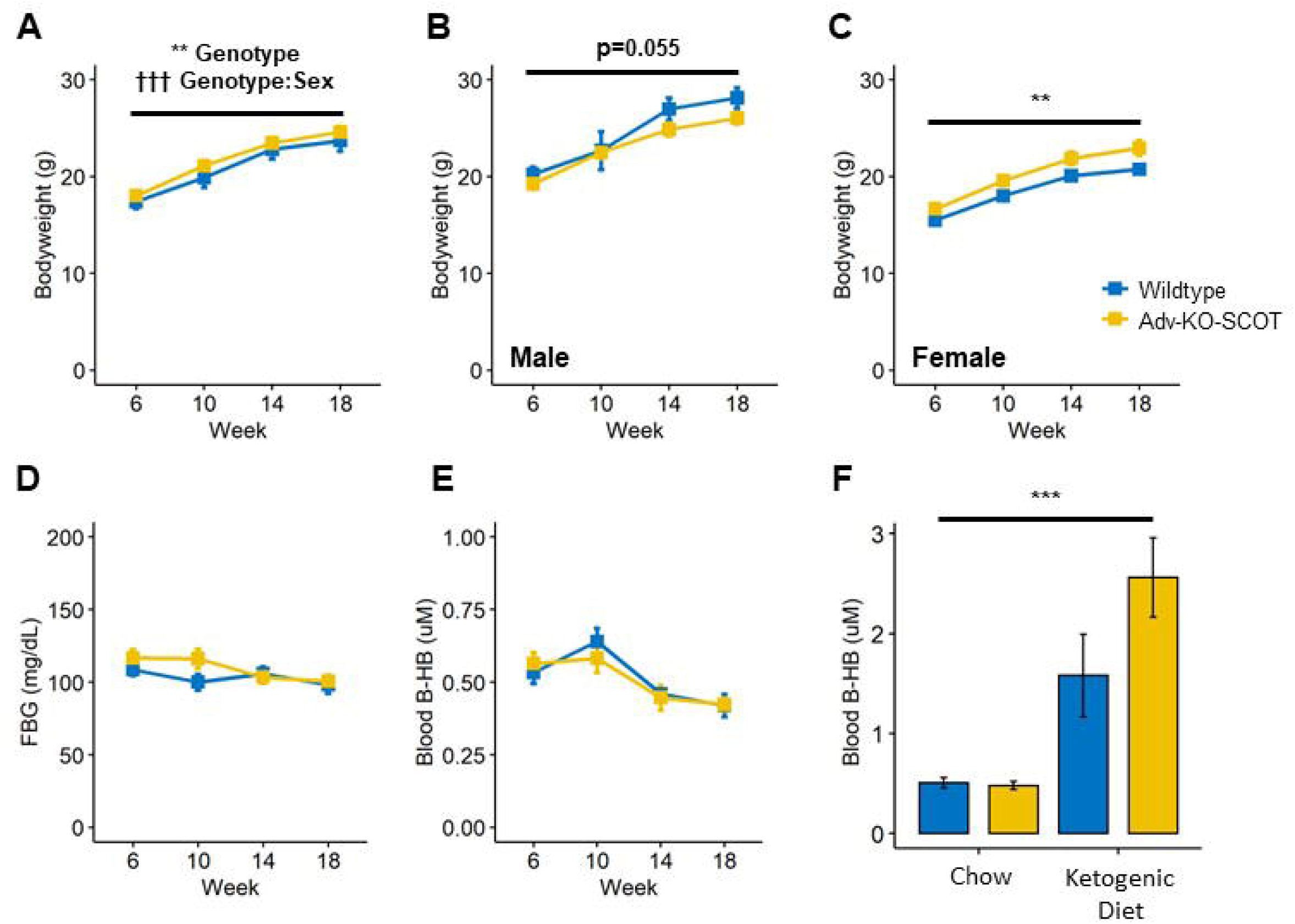
Adv-KO-SCOT mice do not exhibit drastic changes in systemic metabolic markers. (A) The body weight of wildtype and Adv-KO-SCOT mice varied by both genotype and genotype-sex interaction. Male Adv-KO-SCOT mice were largely not different from wildtype mice, though there was a trend toward decreased size (B). Female Adv-KO-SCOT mice, however, had consistently heavier body weight than their littermate, wildtype controls (C). Fasting blood glucose (FBG) (D) and circulating β-hydroxybutyrate (E) were unchanged between Adv-KO-SCOT and wildtype mice. (F) On consumption of a ketogenic diet for one week, circulating β-HB was significantly increased in both genotypes, and there was a trend toward further increased circulating β-HB in Adv-KO-SCOT mice. (D-E) Mice were fasted for three hours before each measure of FBG and β-hydroxybutyrate. N-way, mixed-models ANOVA, ** indicates effect of genotype: p < 0.01, ††† indicates effect of genotype-sex interaction: p < 0.005. (F) N-way ANOVA, *** indicates effect of genotype: p < 0.005.

To assess whether peripheral nerve knockout of SCOT affected circulating ketone levels in the context of ketosis, six-week-old wildtype and Adv-KO-SCOT mice were fed a ketogenic diet for one week. Consumption of a ketogenic diet caused an increase in circulating β-HB in both genotypes (**Figure 2F**; N-way ANOVA, genotype: p = 2.06e^-6^). Additionally, there was a strong but statistically insignificant trend toward an effect of genotype (p = 0.0679) and genotype-diet interaction (p = 0.0775). For all measures of fasting blood glucose and circulating β-HB, mice fasted for 3 hours.

### Adv-KO-SCOT mice have larger TrkA^+^ NF-H^+^ Dorsal Root Ganglia Soma than wildtypes

We harvested DRG from wildtype and Adv-KO-SCOT mice to assess changes in peptidergic (TrkA^+^), non-peptidergic (IB4^+^), and myelinated (neurofilament heavy, NF-H^+^) sensory neuron populations (**Figure 3A**). We performed a Bonferroni correction to account for multiple comparisons in analyzing these populations and set our α-level at 0.005. We detected no significant differences in the area (Student’s t-test, p = 0.332) or diameter (Student’s t-test, p = 0.4761) of all TrkA^+^ soma (**Figure 3B**). Likewise, no changes were present in area (Student’s t-test, p = 0.9375) or diameter (Student’s t-test, p = 0.7313) in IB4^+^ neurons (**Figure 3C**). We detected a statistically significant increase in NF-H^+^ soma area (Student’s t-test, p = 0.002919) but not diameter (Student’s t-test, p = 0.01043) in Adv-KO-SCOT mice (**Figure 3D**). Both area (Student’s t-test, p = 0.001008) and diameter (Student’s t-test, p = 0.0004735) were increased in TrkA^+^ NF-H^+^ DRG soma from Adv-KO-SCOT mice (**Figure 3E**); however, no differences were detected in the area (Student’s t-test, p = 0.6901) or diameter (Student’s t-test, p = 0.5898) of TrkA^+^ NF-H^-^ soma (**Figure 3F**).

**Figure 3.**
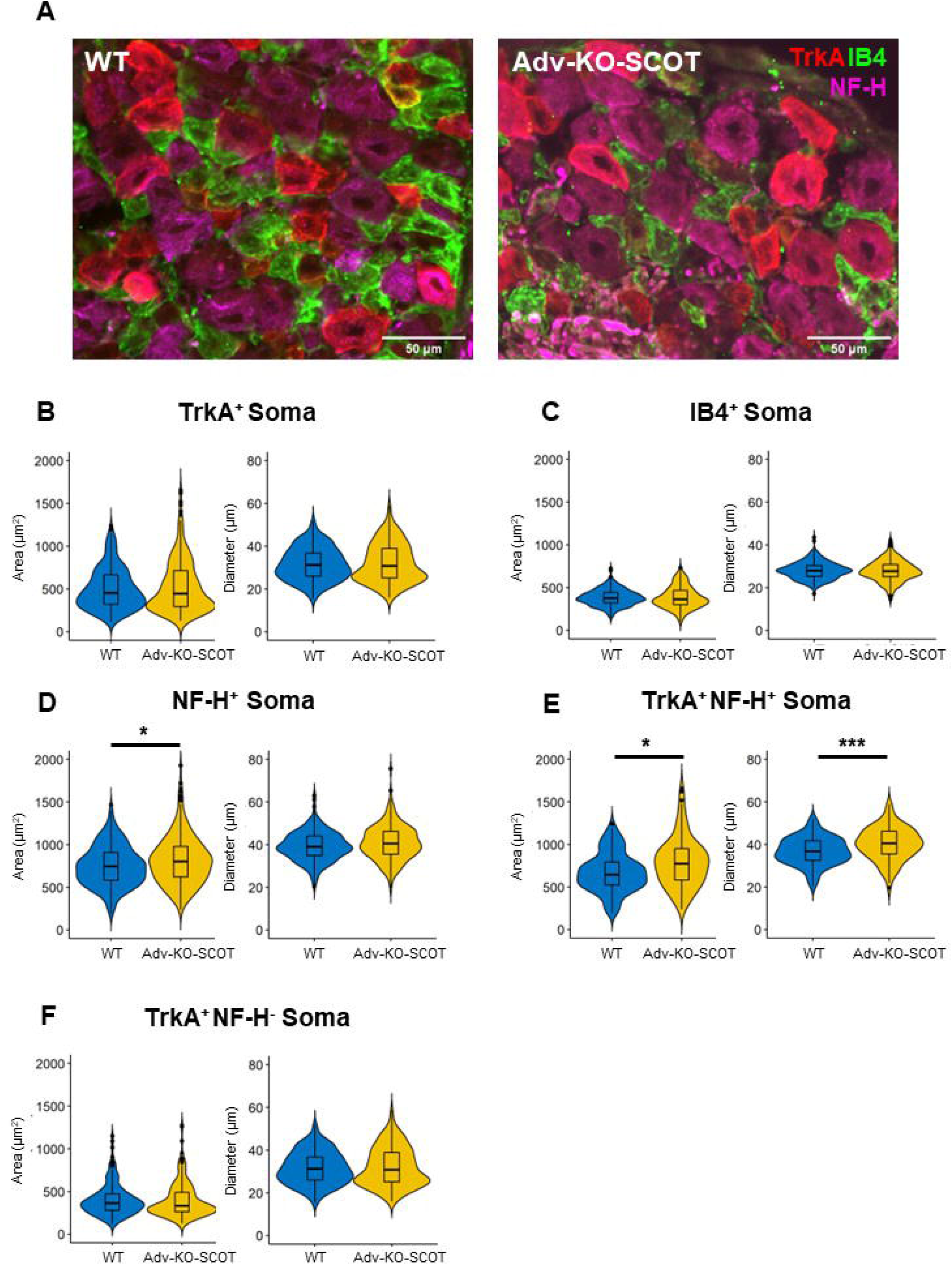
Loss of *Oxct1* during development alters the soma morphology of small, myelinated peptidergic sensory neurons. (A) Representative micrographs of wildtype and Adv-KO-SCOT DRG stained for tropomyosin receptor kinase A (TrkA, red), isolectin B4 (IB4, green), and neurofilament heavy chain (NF-H). Scale bar represents 50 μm. There were no differences in the area or diameter of TrkA^+^ (B) or IB4^+^ (C) soma. There was a significant increase in the area, but not diameter, of NF-H^+^ soma in Adv-KO-SCOT mice (D). Adv-KO-SCOT mice also demonstrated in increased area and diameter of TrkA^+^ NF-H^+^ soma (E), yet there were no differences in the size of TrkA^+^ NF-H^-^ soma (F). Student’s t-test with a Bonferroni correction, * indicates p < 0.005, ** indicates p < 0.001, *** indicates p < 0.0005.

**Figure 4.**
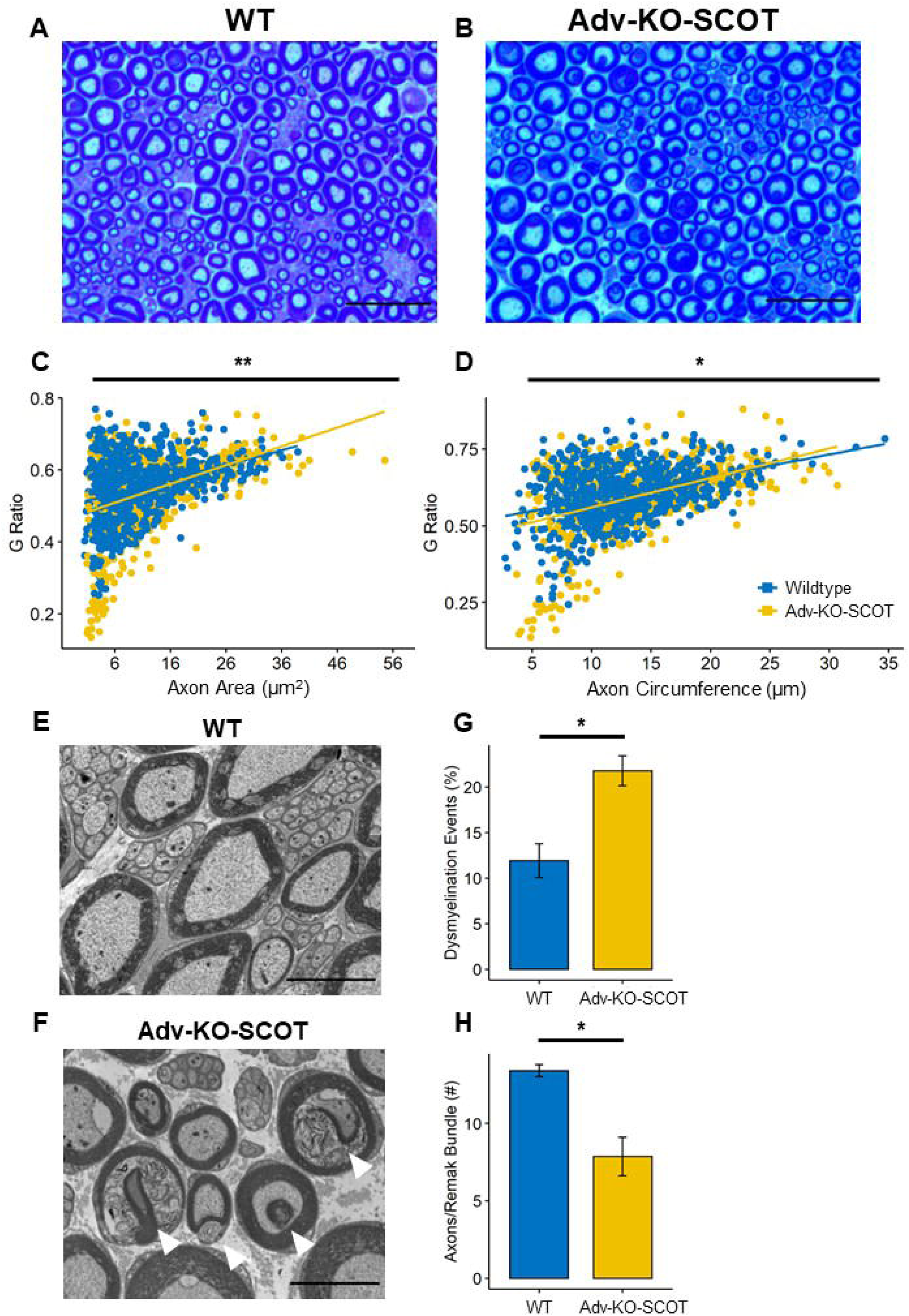
Loss of ketone metabolism in peripheral sensory axons leads to abnormal myelination patterns. (A-B) Representative micrographs of sciatic nerve semi-thin sections stained with toluidine blue used for g-ratio determination (scale bar represents 20 μm). G ratio as determined by myelin and axoplasm area (C) and circumference (D) reveals hypermyelination of small, myelinated axons of the sciatic nerve. Transmission electron micrographs from wildtype (E) and Adv-KO-SCOT (F) mice further revealed myelination errors in Adv-KO-SCOT mice (highlighted by white arrowheads, scale bar represents 5 μm), which are quantified in (G). Additionally, Adv-KO-SCOT mice displayed reduced Remak bundle occupancy compared to wildtype mice (H), indicating both small- and large-fiber deficits in Adv-KO-SCOT mice. (C-D) N-way ANOVA, * indicates effect of genotype: p < 0.05, ** indicates effect of genotype: p < 0.01. (G-H) Student’s t-test, * indicates p < 0.05.

### Myelination aberrations are present in Adv-KO-SCOT mice

We harvested sciatic nerves from 6-week-old Adv-KO-SCOT and wildtype mice and stained 1µm nerve sections in cross-section with toluidine blue to visualize myelin (**Figure 4A-B**). The analysis revealed an abundance of small, hypermyelinated axons in Adv-KO-SCOT mice (**Figure 4B**). These hypermyelinated axons were clearly reflected in g-ratio based on myelin and axon area (Student’s t-test, p = 0.009114) and circumference (Student’s t-test, p = 0.02255) measurements (**Figure 4C-D**). An analysis by transmission electron microscopy similarly revealed an increased in abnormal myelinated axons in Adv-KO-SCOT mice compared to wildtypes (**Figure 4E-G**; Student’s t-test, p = 0.01713). We also detected differences in the number of unmyelinated axons occupying Remak bundles. Remak bundles from wildtype mice contained significantly more C-fibers than Adv-KO-SCOT mice (**Figure 4E-F, H**; Student’s t-test, p = 0.0399).

### Adv-KO-SCOT mice have largely unaltered mechanical and thermal sensitivity

We observed no significant differences in mechanical withdrawal threshold between wildtype and Adv-KO-SCOT mice (**Figure 5A**; N-way mixed-models ANOVA with repeated-measures, genotype: p = 0.645, genotype-time interaction: p = 0.38). Likewise, cold sensitivity was similar between both genotypes as determined by cold plate assay (**Figure 5C**; Student’s t-test, p = 0.2535). Adv-KO-SCOT mice trended toward increased sensitivity to radiant heat (**Figure 5B**; N-way mixed-models ANOVA with repeated-measures, genotype: p = 0.0775, genotype-time interaction: p = 0.0578). Surprisingly, Adv-KO-SCOT mice displayed an attenuated nociceptive response to intraplantar capsaicin injection compared to wildtype mice. Capsaicin injection significantly increased the number of nociceptive events (licking, biting, shaking, etc.) (**Figure 5D**; 2-way ANOVA, genotype: p = 0.023, injection: p = 1.32e^-7^, genotype-injection interaction: p = 0.00502) and time engaged in nociceptive behavior towards the injected paw (**Figure 5E**; 2-way ANOVA, genotype: p = 0.05, injection: p = 2.5e^-5^, genotype-injection interaction: p = 0.0236) in both genotypes. Adv-KO-SCOT mice receiving capsaicin injection engaged in fewer nociceptive events (Tukey’s *post hoc*, Adv-KO-SCOT-capsaicin and wildtype-capsaicin, p = 0.00125) and spent less time engaged in nociceptive behaviors (Tukey’s *post hoc*, Adv-KO-SCOT-capsaicin, and wildtype-capsaicin, p = 0.00987) than wildtype mice receiving capsaicin.

**Figure 5.**
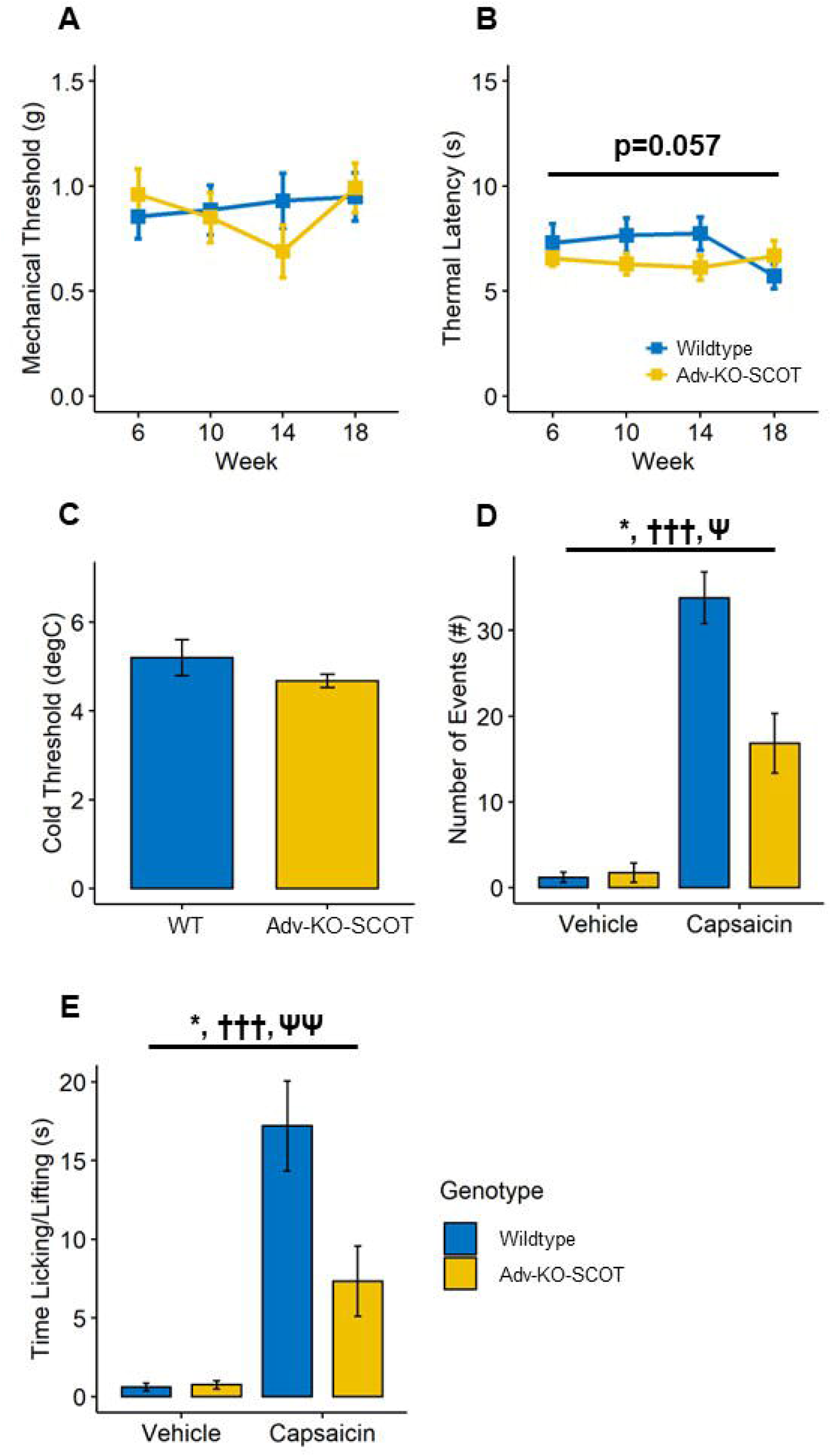
Adv-KO-SCOT mice display normal mechano- and thermal-sensation and hyporesponsiveness to intraplantar capsaicin. (A) Adv-KO-SCOT mice exhibited normal mechanical sensitivity. There was a strong but statistically insignificant trend toward thermal hypersensitivity in Adv-KO-SCOT mice compared to wildtype mice (B). (C) Adv-KO-SCOT and wildtype mice displayed similar sensitivity to cold stimuli as measured by the cold plate test. Adv-KO-SCOT mice engaged in fewer nocifensive events (licking, biting, lifting, etc.) (D) and spent less time engaged in nocifensive behaviors (E) in response to intraplantar capsaicin compared to wildtype mice. (A-B) N-way, mixed models ANOVA. (D-E) N-way ANOVA, * indicates main effect of genotype: p < 0.05, ††† indicates main effect of capsaicin: p < 0.005, Ψ indicates interaction effect of genotype-capsaicin: p < 0.05, ΨΨ indicates interaction effect of genotype-capsaicin: p < 0.01.

### Adv-KO-SCOT mice have decreased epidermal innervation

To assess whether absent ketolysis in peripheral neurons affected epidermal innervation, we harvested footpads from wildtype and Adv-KO-SCOT mice at 6 weeks of age and used PGP9.5 and tropomyosin receptor kinase A (TrkA) antibodies to visualize axonal populations (**Figure 6A**). Using these antibodies, PGP9.5^+^TrkA^+^ axons were considered peptidergic, while PGP9.5^+^TrkA^-^ axons were counted as nonpeptidergic epidermal axons. Adv-KO-SCOT mice had a significantly reduced overall intraepidermal nerve fiber (IENF) density (**Figure 6B**; Student’s t-test, p = 0.0042), reduced peptidergic IENF density (**Figure 6C**; Student’s t-test, p = 0.00279), and reduced nonpeptidergic IENF density (**Figure 6D**, Student’s t-test, p = 0.014). Adv-KO-SCOT mice had a slight but statistically significant increased ratio of nonpeptidergic to peptidergic IENFs compared to wildtype mice (**Figure 6E**; Student’s t-test, p = 0.0455).

**Figure 6.**
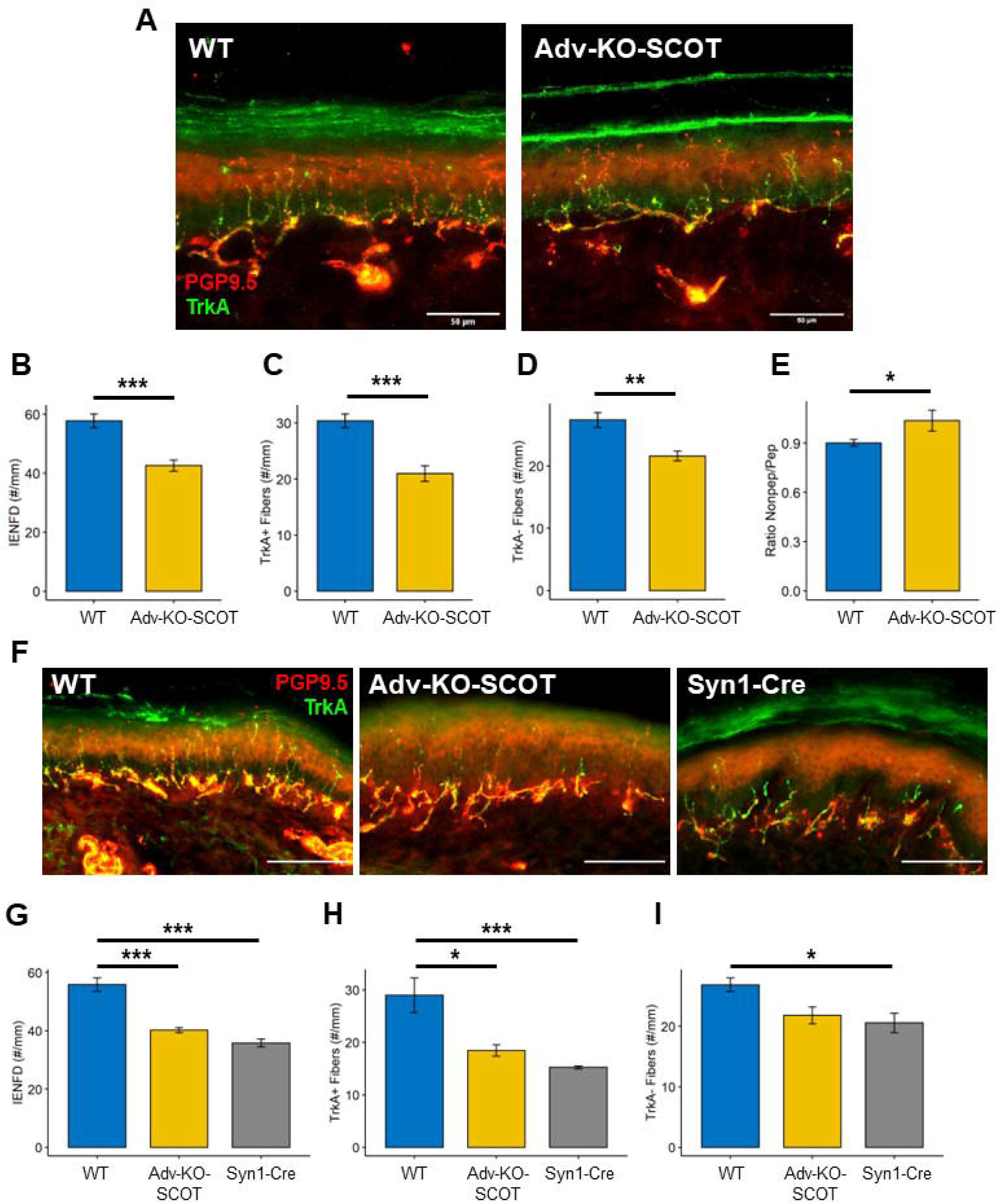
Loss of neuronal ketone metabolism decreases intraepidermal nerve fiber density. (A) Representative micrographs of intraepidermal nerve fibers (IENFs) stained with post-gene product 9.5 (PGP9.5; red) and tropomyosin-receptor kinase A (TrkA, green) from glabrous skin of the hindpaw from 6-week-old wildtype and Adv-KO-SCOT mice (scale bar represents 50 μm). Adv-KO-SCOT mice had a lower total IENF density compared to wildtype mice (B). Adv-KO-SCOT mice exhibited a decrease in both TrkA^+^, peptidergic IENF density (C) and TrkA^-^, nonpeptidergic IENF density (D). There was a slight but statistically significant increase in nonpeptidergic to peptidergic IENF density ratio in Adv-KO-SCOT mice compared to wildtype mice (E). (F) Representative micrographs of IENFs stained with PGP9.5 (red) and TrkA (green) from the glabrous skin of the hindpaw from 6–7-month-old wildtype, Adv-KO-SCOT, and Synapsin-1^+^/^Cre^*Oxct1^flox/flox^*(Syn1-Cre) mice (scale bar represents 100 μm). There was a significant effect of genotype on IENF density for total, peptidergic, and nonpeptidergic fibers (G-I). Adv-KO-SCOT and Syn1-Cre mice had lower total (G) and peptidergic (H) IENF density than wildtype controls. (I) Only Syn1-Cre mice had significantly reduced nonpeptidergic IENF density compared to wildtype controls in adult mice. (B-E) Student’s t-test, * indicates p < 0.05, ** indicates p < 0.01, *** indicates p < 0.005. (G-I) 1-way ANOVA, with a statistically significant effect of genotype in all figures. Tukey’s *post hoc* test, * indicates p < 0.05 for the comparison indicated by black line, *** indicates p < 0.005 for the comparison indicated by black line.

We obtained footpad tissue from an alternative mouse line in which Oxct1 was knocked out using a pan-neuronal promoter (*Syn1^+/Cre^Oxct1^flox/flox^* mice, hereafter Syn1-Cre). Comparisons of epidermal innervation of 26-30-week-old wildtype, Adv-KO-SCOT, and pan-neuronal (Syn1-Cre) mice (**Figure 6F)** revealed significantly reduced IENF density in both Adv-KO-SCOT and Syn1-Cre mice (**Figure 6G**; 1-way ANOVA, genotype: p = 9.5e^-5^; Tukey’s *post hoc*, wildtype-Adv-KO-SCOT: p = 0.000342, wildtype-Syn1-Cre: p = 0.000106). Both Adv-KO-SCOT and Syn1-Cre mice had significantly fewer PGP9.5^+^TrkA^+^ peptidergic fibers compared to wildtype mice (**Figure 6H**, 1-way ANOVA, genotype: p = 0.00362; Tukey’s *post hoc*, wildtype-Adv-KO-SCOT: p = 0.011, wildtype-Syn1-Cre: p = 0.0039). Only Syn1-Cre had significantly reduced TrkA-non-peptidergic IENF density compared to wildtype mice (**Figure 6I**, 1-way ANOVA, genotype: p = 0.0427; Tukey’s *post hoc*, wildtype-Adv-KO-SCOT: p = 0.0847, wildtype-Syn1-Cre: p = 0.0471).

### Adv-KO-SCOT mice have abnormal afferent innervation of the spinal dorsal horn

We assessed the effect of the absence of sensory axonal ketolysis on sensory axon terminations in the dorsal horn of the lumbar spinal cord (**Figure 7A**). We detected no difference in the distribution of TrkA^+^ fibers in the superficial Rexed lamina I between Adv-KO-SCOT and wildtype mice (**Figure 7B**; Student’s t-test, p = 0.8394). Likewise, we detected no difference in IB4^+^ innervation of lamina IIo (**Figure 7C**; Student’s t-test, p = 0.8207). We did, however, notice more TrkA^+^ projections extending into the deeper lamina of the dorsal horn in Adv-KO-SCOT mice compared to WT (**Figure 7A**, *white arrowheads*). Following Bonferroni corrections for multiple comparisons (n comparisons = 3), counts of the number of TrkA^+^ axons extending to deeper lamina reached statistical significance (**Figure 7D**, Student’s t-test, p = 0.01551).

**Figure 7.**
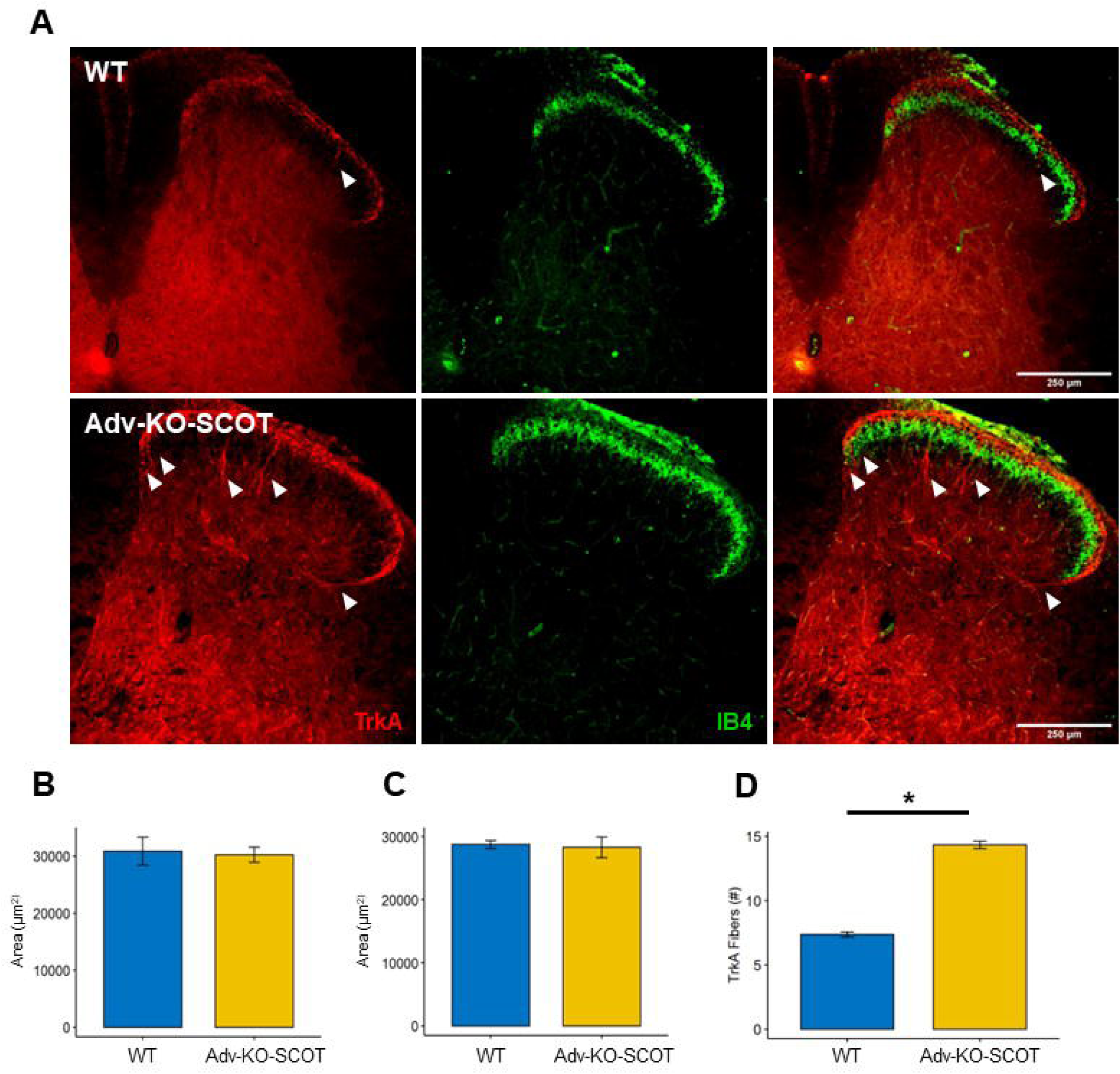
*Oxct1* deficiency results in abnormal peripheral sensory innervation of the spinal dorsal horn. (A) Representative micrographs of lumbar spinal dorsal horns from 6-week-old Adv-KO-SCOT and wildtype mice stained with TrkA (red) and isolectin B4 (IB4, green; scale bar represents 250 μm). There were no differences in the areas of peptidergic (TrkA, B) or nonpeptidergic (IB4, C) innervation of superficial lamina from Adv-KO-SCOT and wildtype mice. The spinal dorsal horn from Adv-KO-SCOT mice did contain more TrkA^+^ fibers penetrating through the superficial laminae into deeper areas for the spinal dorsal horn (quantified in D; marked by white arrowheads in A). Student’s t-test, * indicates p < 0.05.

### Adv-KO-SCOT mice exhibit impaired proprioception and coordination

We tested gross coordination and balance in male and female Adv-KO-SCOT and wildtype mice using a Rotorod. We detected no significant differences in latency to fall between Adv-KO-SCOT and wildtype mice (**Figure 8A**, 2-way ANOVA, genotype: p = 0.549, sex: p = 0.0696, genotype-sex interaction: p = 0.3092). As an additional measure of proprioception, mice were subjected to a grid-walk test. During a timed grid-walk, Adv-KO-SCOT mice spent less time walking on the grid than wildtype mice (**Figure 8B** 2-way ANOVA, genotype: p = 2.43e^-5^, sex: p = 0.639, genotype-sex interaction: p = 0.909). When normalized to the amount of time walking, Adv-KO-SCOT mice also exhibited an increased number of total paw slips (**Figure 8C**, 2-way ANOVA, genotype: p = 1.03e^-10^, sex: p = 0.401, genotype-sex interaction: p = 0.234), which was attributed to impaired coordination of both the forepaw (**Figure 8D**, 2-way ANOVA, genotype: p = 2.79e^-10^, sex: p = 0.373, genotype-sex interaction: p = 0.122) and hind paw (**Figure 8E**, 2-way ANOVA, genotype: p = 4.51e^-8^, sex: p = 0.657, genotype-sex interaction: p = 0.783).

**Figure 8.**
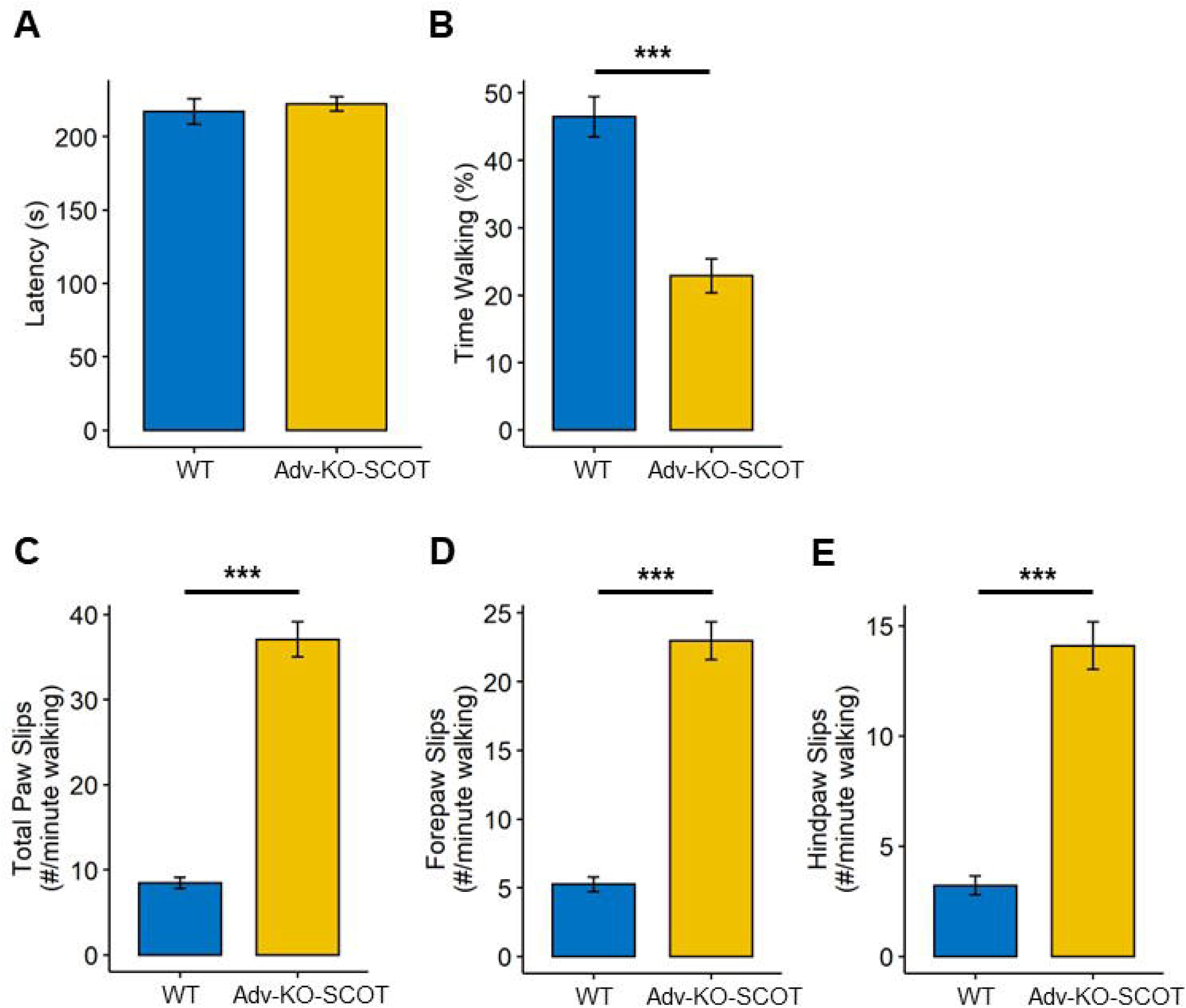
Loss of ketone metabolism in peripheral sensory axons results in proprioceptive deficits. (A) Adv-KO-SCOT mice did not display any differences compared to wildtype mice in latency to fall during the rotarod test. During grid-walk test, Adv-KO-SCOT mice spent significantly less time walking than wildtype mice (B). Additionally, Adv-KO-SCOT mice exhibited a significant increase in total paw slips normalized to time walking during gridwalk (C). These effects were not limb dependent, as Adv-KO-SCOT mice exhibit more forepaw slips (D) and hindpaw slips (E) normalized to time compared to wildtype mice. N-way ANOVA, *** indicates a significant effect of genotype: p < 0.005.

### Adv-KO-SCOT mice have normal innervation of the muscle spindle

We assessed sensory innervation of the muscle in Adv-KO-SCOT by staining the muscle spindles of the gastrocnemius. Adv-KO-SCOT mice did not display any obvious denervation or structural deficits in the muscle spindle (**Figure 9A**). Wildtype and Adv-KO-SCOT mice did not have differences in the distance between axonal rotations (**Figure 9B**; Student’s t-test, p = 0.172) that were statistically significant. Likewise, the width of NEF-H^+^ axonal rotations at the muscle spindle was not different between Adv-KO-SCOT and wildtype mice (**Figure 9C**; Student’s t-test, p = 0.387).

**Figure 9.**
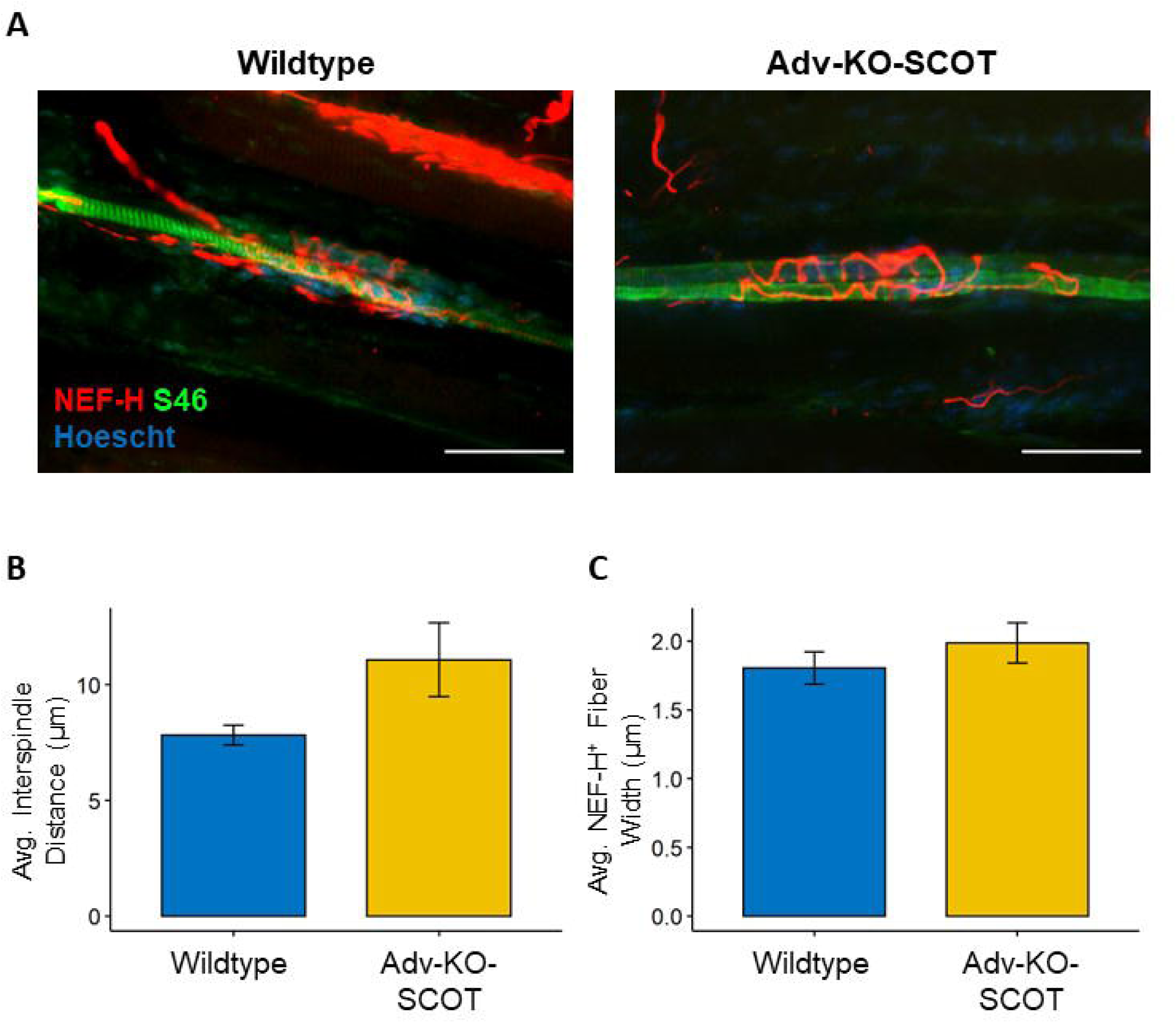
SCOT knockout does not affect innervation of the muscle spindle. (A) Representative micrographs of innervation of the muscle spindle in the gastrocnemius from six-week-old wildtype and Adv-KO-SCOT mice stained with NEF-H (red), S46 (green), and Hoescht (blue; scale bar represents 50 μm). There were no significant differences in the interspindle distance between NEF-H^+^ axonal rotations (B) or in the width of the axon in those rotations (C) between wildtype and Adv-KO-SCOT mice. (B-C) Student’s t-test.

## Discussion

There is mounting evidence that ketone utilization is protective (Cooper et al., 2018b, Ruskin et al., 2021, Zhong et al., 2021) and, in some cases, promotes the regeneration (Enders et al., 2022a) in the somatosensory nervous system of rodent models of chronic pain and neuropathy. SCOT is a mitochondrial enzyme expressed in peripheral tissues and required for ketolysis, catalyzing the conversion of acetoacetate and succinyl-CoA to acetoacetyl-CoA and succinate. *Oxct1* mRNA and SCOT protein are downregulated in rodent models of amyotrophic lateral sclerosis (Szelechowski et al., 2018) and Friedreich’s ataxia (Dong et al., 2022), as well as in fibroblasts from Friedreich’s ataxia patients (Dong et al., 2022). The contribution of impaired ketone metabolism in these diseases is unclear. In addition, relatively little is known about the importance of ketone metabolism in the normal development and maintenance of the somatosensory nervous system.

We generated tissue-specific knockouts of SCOT in the peripheral nervous system of mice using advillin-driven expression of Cre recombinase. Wildtype mice expressed SCOT in the soma of DRG neurons, while we were unable to detect SCOT expression in Adv-KO-SCOT DRG. Existing single cell-transcriptomics datasets from mouse and human suggest that *Oxct1* is expressed in virtually all described sensory neuronal populations in mouse DRG (Usoskin et al., 2015, Sharma et al., 2020) and putative peptidergic C-fibers, Aδ-nociceptors, and Aδ-LTMRs in human DRG (Nguyen et al., 2021). GFAP^+^ cells in the DRG of wildtype and Adv-KO-SCOT mice did not express SCOT, indicating that these cells are likely unable to oxidize ketones.

We did not detect changes in the concentrations of circulating fuel sources (e.g., fasting blood glucose, β-HB) between chow-fed wildtype and Adv-KO-SCOT mice. However, while circulating ketones were elevated in both genotypes following administration of a ketogenic diet, Adv-KO-SCOT mice had significantly increased blood ketones compared to wildtype mice. Ketone bodies are the primary circulating fuel source during nutritional ketosis. Thus, increased accumulation of blood ketones in mice with deficient ketolysis in the somatosensory nervous system is indicative of the high metabolic demands of the peripheral nervous system.

Our studies demonstrate that sensory neuron-specific knockout of *Oxct1* leads to pathological changes in the somatosensory nervous system. Adv-KO-SCOT mice have significantly fewer IENFs in the glabrous skin of the hindpaw. IENFs are comprised of unmyelinated C-fibers, which, in mice, can be subdivided into developmentally distinct peptidergic and non-peptidergic axon populations (Molliver et al., 1997), marked by the presence or absence of TrkA and positive staining for neuropeptides or the lectin IB4, respectively. These epidermal fibers degenerate at different rates during diabetes (Johnson et al., 2008) and following nerve injury (Wang et al., 2011), and changes in the proportion of “non-peptidergic” to “peptidergic” fibers have been associated with abnormal sensation in obese mice (Groover et al., 2013). Six-week-old Adv-KO-SCOT mice exhibited reduced cutaneous innervation by both “peptidergic” (PGP9.5^+^ TrkA^+^,) and “non-peptidergic” (PGP9.5^+^ TrkA^-^,) fibers. While Adv-KO-SCOT mice had fewer IENFs than wildtype mice, we did not detect any differences in “peptidergic” (TrkA^+^ NF-H^-^) or “non-peptidergic” (IB4^+^) soma in the DRG between genotypes. However, the consistency of epidermal axon loss in Adv-KO-SCOT and Syn1-Cre Oxct1 mice suggests that axonal ketone oxidation is critical to developing and maintaining a full complement of axonal terminations in the epidermis.

Loss of IENF density is a hallmark of small-fiber neuropathy and is observed in various conditions associated with chronic pain in both humans (Chien et al., 2020, Cheng et al., 2013, Bednarik et al., 2009, Kluding et al., 2012) and rodents (Jack et al., 2012, Johnson et al., 2008, Cooper et al., 2018b, Enders et al., 2022a). Prior work from our laboratory has demonstrated that sensory abnormalities usually precede these anatomic changes (Cooper et al., 2018b, Johnson et al., 2008). While Adv-KO-SCOT mice exhibit lower IENF density than wildtype mice, their mechanical and thermal thresholds remained largely unchanged. Adv-KO-SCOT mice demonstrated no significant changes in responsiveness to mechanical or cold stimuli. There was a strong trend toward increased responsiveness to radiant heat in Adv-KO-SCOT mice that was lost with age, though this failed to reach statistical significance. The discrepancy between reduced epidermal innervation and normal sensory thresholds is consistent with the view that these two aspects are not always linked (Cooper et al., 2018b, Wright et al., 2007, Tierney et al., 2022). Surprisingly, Adv-KO-SCOT mice exhibited fewer nocifensive behaviors following capsaicin injection than their wildtype littermates. Thus, our results suggest that deficiencies in ketone oxidation in peripheral sensory neurons may be more important in responses to noxious chemicals like capsaicin.

In addition to altered cutaneous innervation, loss of *Oxct1* was associated with increased sizes of myelinated, peptidergic DRG soma. This neuronal population fits the PEP2 classification (TrkA^+^ NF-H^+^), representing nociceptive Aδ fibers, described by Usoskin et. al. (Usoskin et al., 2015). However, single-cell quantitative polymerase chain reaction and electrophysiological profiling of the DRG suggests that TrkA^+^ NF-H^+^ soma represent A-fiber high-threshold mechanoreceptors (HTMRs) or another population of A-fibers (Adelman et al., 2019). Adelman and colleagues describe this other population as large dynamic range neurons, though their characteristics are consistent with low-threshold mechanoreceptors (LTMRs)(Adelman et al., 2019). Our analysis was insufficient to determine whether the increased size in TrkA^+^ NF-H^+^ soma represents a change in either HTMRs or LTMRs, as Adv-KO-SCOT mice exhibited no changes in mechanical sensitivity. Electrophysiological profiling of these mice is thus required to confirm the functional identity of this population, though both studies above indicate these neurons likely represent Aδ sensory neurons.

The changes in TrkA^+^ NF-H^+^ neurons were accompanied by abnormal myelination of axons in the sciatic nerve of Adv-KO-SCOT mice. Adv-KO-SCOT mice demonstrated a clear tail of decreased g-ratio in axons smaller than 10 μm^2^, indicating that these smaller fibers are hypermyelinated. This area correlates to ∼1.8 μm diameter axons, well within the fiber diameters of putative Aδ neurons. It is unclear why Aδ neurons would be uniquely susceptible during development without ketone oxidation compared to unaffected populations. Aδ HTMRs are critically reliant on nerve growth factor (NGF)-TrkA signaling between postnatal day 0 and 14 (Lewin et al., 1992), and advillin expression—and thus, *Oxct1* knockout in our animals—is present in the DRG by postnatal day 0 (Zurborg et al., 2011). Ketone metabolism may dampen the NGF-TrkA signaling axis. This explanation is consistent with somal and myelination changes in an Aδ population, as NGF signaling promotes myelination in the peripheral nervous system (Chan et al., 2004).

It is also possible that decreased axonal ketone oxidation increases the availability of ketone bodies for myelin biosynthesis. Radiolabeling studies demonstrate the preferential use of ketone-derived acetyl groups in myelin sterol biosynthesis in the rat brain (Koper et al., 1981) and mouse sciatic nerve (Clouet and Bourre, 1988) over those derived from glycolysis. As sensory axons no longer consume ketones as fuel in these mice, myelinating Schwann cells may instead shuttle β-HB and acetoacetate into myelin lipid synthesis, explaining the hypermyelination phenotype. While chow-fed Adv-KO-SCOT mice do not exhibit elevated circulating β-HB levels compared to wildtype mice, there was a strong trend toward increased circulating β-HB levels in ketogenic diet-fed Adv-KO-SCOT mice compared to wildtype (**Figure 2**). This indicates there is no change in basal ketone utilization even in the absence of axonal ketolysis. These data are consistent with alternative avenues of ketone utilization, including incorporation into myelin cholesterol. Further, the trend toward increased circulating blood ketones in Adv-KO-SCOT mice during nutritional ketosis suggests Adv-KO-SCOT mice lose SCOT function in the somatosensory nervous system.

In addition to sensory changes, myelination pathology, and axon deficiencies and abnormalities, loss of *Oxct1* in sensory neurons led to gait dysfunction. An analysis of grid-walk revealed significant disruptions to fore- and hind-limb proprioception in the Adv-KO-SCOT mice. This pattern of sensory deficits is relevant, as a recent study has reported that *Oxct1* expression is downregulated in both mouse models and patient samples of Friedreich’s ataxia (Dong et al., 2022). The neurological symptoms of Friedreich’s ataxia include a loss of cutaneous sensation, altered gait, fatigue, and worsening balance and proprioception (Creigh et al., 2019, Delatycki et al., 2012, Nolano et al., 2001). Mouse models of Friedreich’s ataxia likewise demonstrate altered gait, poor proprioception, and altered innervation of the muscle spindle (Gérard et al., 2023, Piguet et al., 2018, Mollá et al., 2016). In this new study, mutant frataxin leads to increased ubiquitination of SCOT, leading to its degradation and severely impaired ketone metabolism in fibroblasts and skeletal muscle (Dong et al., 2022). While Adv-KO-SCOT mice did not recapitulate abnormal innervation of muscle spindles as seen in the YG8R mouse model of Friedreich’s ataxia (Mollá et al., 2016), this discrepancy may be due to age. Others have described abnormal width of axonal ensheathment of the muscle spindle in aged, 24-month-old mice, while we assessed innervation of the muscle spindle in 6-week-old mice. As our analysis of Adv-KO-SCOT mice was in relatively young mice, it will be important to determine the breadth of sensory changes as these mice age and how these changes correlate with the breadth of symptoms in patients with Friedreich’s ataxia (Creigh et al., 2019, Nolano et al., 2001). Our work suggests that impaired ketone metabolism in peripheral sensory neurons can mimic the neurological deficits of Friedreich’s ataxia and supports the model in which mutant frataxin mediated downregulates SCOT and ketolysis, contributing to the clinical symptoms of this disease.

The importance of proper energy metabolism in the function of the somatosensory nervous system is emerging rapidly. Neurodegenerative diseases have recently revealed deficiencies in ketolysis, including Friedreich’s ataxia and amyotrophic lateral sclerosis (Dong et al., 2022, Szelechowski et al., 2018). Meanwhile, ketogenic diets are being studied in various neuropathies (Cooper et al., 2018b, Enders et al., 2022b, Field et al., 2022, Ruskin et al., 2009, Ruskin et al., 2021, Zhong et al., 2021, Enders et al., 2022a). Our findings on the loss of *Oxct1* and SCOT in peripheral neurons provide new information toward our understanding of ketone oxidation for developing and maintaining the peripheral sensory nervous system. Overall, these findings suggest that ketone oxidation is essential for the normal development of sensory axons, highlights the deleterious effects of deficient ketone metabolism during development and provides a new link to possible mechanisms leading to sensory dysfunction in patients with Friedreich’s ataxia.

## Acknowledgments

This work was supported by NIH grants R01 NS043314 (DEW), R01 AG069781 (PAC and JPT), R01 DK091538 (PAC), the Kansas Institutional Development Award (IDeA) P20 GM103418, Kansas University Training Program in Neurological and Rehabilitation Sciences NIH T32HD057850 (JE), Translating Obesity, Metabolic Dysfunction, and Comorbid Disease States NIH T32DK128770 (ST), the Kansas IDDRC P30HD00228, and University of Minnesota Institute for Diabetes, Obesity, and Metabolism.

## Author contributions

JE and DEW designed the research study; JE, JMR, MTS, PL, JJ, ST, SL, and XC performed the experiments; JE and DEW analyzed the data; all authors contributed to the manuscript.

## Bibliography

Adelman, P. C., Baumbauer, K. M., Friedman, R., Shah, M., Wright, M., Young, E., Jankowski, M. P., Albers, K. M. & Koerber, H. R. 2019. Single-cell q-Pcr derived expression profiles of identified sensory neurons. Molecular Pain, 15, 1744806919884496.

Bednarik, J., Vlckova-Moravcova, E., Bursova, S., Belobradkova, J., Dusek, L. & Sommer, C. 2009. Etiology of small-fiber neuropathy. Journal of the Peripheral Nervous System, 14, 177–183.

Bilger, A. & Nehlig, A. 1992. Quantitative histochemical changes in enzymes involved in energy metabolism in the rat brain during postnatal development. Ii. Glucose-6-phosphate dehydrogenase and β-hydroxybutyrate dehydrogenase. International Journal of Developmental Neuroscience, 10, 143–152.

Bongiovanni, D., Benedetto, C., Corvisieri, S., Del Favero, C., Orlandi, F., Allais, G., Sinigaglia, S. & Fadda, M. 2021. Effectiveness of ketogenic diet in treatment of patients with refractory chronic migraine. Neurological Sciences, 42, 3865–3870.

Chan, J. R., Watkins, T. A., Cosgaya, J. M., Zhang, C., Chen, L., Reichardt, L. F., Shooter, E. M. & Barres, B. A. 2004. Ngf Controls Axonal Receptivity to Myelination by Schwann Cells or Oligodendrocytes. Neuron, 43, 183–191.

Chaplan, S. R., Bach, F. W., Pogrel, J. W., Chung, J. M. & Yaksh, T. L. 1994. Quantitative assessment of tactile allodynia in the rat paw. J Neurosci Methods, 53, 55–63.

Cheng, H. T., Dauch, J. R., Porzio, M. T., Yanik, B. M., Hsieh, W., Smith, A. G., Singleton, J. R. & Feldman, E. L. 2013. Increased axonal regeneration and swellings in intraepidermal nerve fibers characterize painful phenotypes of diabetic neuropathy. The journal of pain, 14, 941–947.

Chien, Y.-L., Chao, C.-C., Wu, S.-W., Hsueh, H.-W., Chiu, Y.-N., Tsai, W.-C., Gau, S. S.-F. & Hsieh, S.-T. 2020. Small fiber pathology in autism and clinical implications. Neurology, 95, e2697–e2706.

Clouet, P. M. & Bourre, J.-M. 1988. Ketone Body Utilization for Lipid Synthesis in the Murine Sciatic Nerve: Alterations in the Dysmyelinating Trembler Mutant. Journal of Neurochemistry, 50, 1494–1497.

Cooper, M. A., Mccoin, C., Pei, D., Thyfault, J. P., Koestler, D. & Wright, D. E. 2018a. Reduced mitochondrial reactive oxygen species production in peripheral nerves of mice fed a ketogenic diet. Experimental physiology, 103, 1206–1212.

Cooper, M. A., Menta, B. W., Perez-Sanchez, C., Jack, M. M., Khan, Z. W., Ryals, J. M., Winter, M. & Wright, D. E. 2018b. A ketogenic diet reduces metabolic syndrome-induced allodynia and promotes peripheral nerve growth in mice. Experimental Neurology, 306, 149–157.

Cotter, D. G., Schugar, R. C., Wentz, A. E., André D’avignon, D. & Crawford, P. A. 2013. Successful adaptation to ketosis by mice with tissue-specific deficiency of ketone body oxidation. American Journal of Physiology-Endocrinology and Metabolism, 304, E363–E374.

Creigh, P. D., Mountain, J., Sowden, J. E., Eichinger, K., Ravina, B., Larkindale, J. & Herrmann, D. N. 2019. Measuring peripheral nerve involvement in Friedreich’s ataxia. Annals of clinical and translational neurology, 6, 1718–1727.

Delatycki, M. B., Corben, L. A. & Maria, B. L. 2012. Clinical Features of Friedreich Ataxia. Journal of Child Neurology, 27, 1133–1137.

Di Lorenzo, C., Coppola, G., Bracaglia, M., Di Lenola, D., Sirianni, G., Rossi, P., Di Lorenzo, G., Parisi, V., Serrao, M. & Cervenka, M. C. 2019. A ketogenic diet normalizes interictal cortical but not subcortical responsivity in migraineurs. Bmc neurology, 19, 1–9.

Dong, Y. N., Mesaros, C., Xu, P., Mercado-Ayon, E., Halawani, S., Ngaba, L., Warren, N., Sleiman, P., Rodden, L. & Schadt, K. A. 2022. Frataxin controls ketone body metabolism through regulation of OXCT1. Pnas Nexus.

Enders, J., Swanson, M. T., Ryals, J. & Wright, D. E. 2022a. A ketogenic diet reduces mechanical allodynia and improves epidermal innervation in diabetic mice. Pain, 163, 682–689.

Enders, J., Thomas, S., Swanson, M. T., Ryals, J. M. & Wright, D. E. 2022b. A ketogenic diet prevents methylglyoxal-evoked nociception by scavenging methylglyoxal. The Journal of Pain, 23, 24.

Field, R., Pourkazemi, F. & Rooney, K. 2022. Effects of a Low-Carbohydrate Ketogenic Diet on Reported Pain, Blood Biomarkers and Quality of Life in Patients with Chronic Pain: A Pilot Randomized Clinical Trial. Pain Medicine, 23, 326–338.

Gérard, C., Archambault, A. F., Bouchard, C. & Tremblay, J. P. 2023. A promising mouse model for Friedreich Ataxia progressing like human patients. Behavioural Brain Research, 436, 114107.

Groover, A. L., Ryals, J. M., Guilford, B. L., Wilson, N. M., Christianson, J. A. & Wright, D. E. 2013. Exercise-mediated improvements in painful neuropathy associated with prediabetes in mice. Pain®, 154, 2658–2667.

Henderson, C. B., Filloux, F. M., Alder, S. C., Lyon, J. L. & Caplin, D. A. 2006. Efficacy of the ketogenic diet as a treatment option for epilepsy: meta-analysis. Journal of child neurology, 21, 193–198.

Hugo, S. E., Cruz-Garcia, L., Karanth, S., Anderson, R. M., Stainier, D. Y. & Schlegel, A. 2012. A monocarboxylate transporter required for hepatocyte secretion of ketone bodies during fasting. Genes Dev, 26, 282–93.

Izumi, Y., Ishii, K., Katsuki, H., Benz, A. M. & Zorumski, C. F. 1998. beta-Hydroxybutyrate fuels synaptic function during development. Histological and physiological evidence in rat hippocampal slices. The Journal of clinical investigation, 101, 1121–1132.

Jack, M. M., Ryals, J. M. & Wright, D. E. 2012. Protection from diabetes-induced peripheral sensory neuropathy — A role for elevated glyoxalase I? Experimental Neurology, 234, 62–69.

Johnson, M. S., Ryals, J. M. & Wright, D. E. 2008. Early loss of peptidergic intraepidermal nerve fibers in an Stz-induced mouse model of insensate diabetic neuropathy. Pain, 140, 35–47.

Kluding, P. M., Pasnoor, M., Singh, R., Jernigan, S., Farmer, K., Rucker, J., Sharma, N. K. & Wright, D. E. 2012. The effect of exercise on neuropathic symptoms, nerve function, and cutaneous innervation in people with diabetic peripheral neuropathy. Journal of Diabetes and its Complications, 26, 424–429.

Koper, J. W., Lopes-Cardozo, M. & Van Golde, L. M. G. 1981. Preferential utilization of ketone bodies for the synthesis of myelin cholesterol in vivo. Biochimica et Biophysica Acta (Bba) - Lipids and Lipid Metabolism, 666, 411–417.

Kraus, H., Schlenker, S. & Schwedesky, D. 1974. Developmental Changes of Cerebral Ketone Body Utilization in Human Infants. Biological Chemistry, 355, 164–170.

Lewin, G. R., Ritter, A. M. & Mendell, L. 1992. On the role of nerve growth factor in the development of myelinated nociceptors. Journal of Neuroscience, 12, 1896–1905.

Massieu, L., Haces, M., Montiel, T. & Hernandez-Fonseca, K. 2003. Acetoacetate protects hippocampal neurons against glutamate-mediated neuronal damage during glycolysis inhibition. Neuroscience, 120, 365–378.

Mollá, B., Riveiro, F., Bolinches-Amorós, A., Muñoz-Lasso, D. C., Palau, F. & González-Cabo, P. 2016. Two different pathogenic mechanisms, dying-back axonal neuropathy and pancreatic senescence, are present in the YG8r mouse model of Friedreich’s ataxia. Disease Models & Mechanisms, 9, 647–657.

Molliver, D., Wright, D., Leitner, M., Parsadanian, A. S., Doster, K., Wen, D., Yan, Q. & Snider, W. 1997. IB4-binding Drg neurons switch from Ngf to Gdnf dependence in early postnatal life. Neuron, 19, 849–861.

Nehlig, A., Boyet, S. & De Vasconcelos, A. P. 1991. Autoradiographic measurement of local cerebral β-hydroxybutyrate uptake in the rat during postnatal development. Neuroscience, 40, 871–878.

Nguyen, M. Q., Von Buchholtz, L. J., Reker, A. N., Ryba, N. J. P. & Davidson, S. 2021. Single-nucleus transcriptomic analysis of human dorsal root ganglion neurons. eLife, 10, e71752.

Nolano, M., Provitera, V., Crisci, C., Saltalamacchia, A. M., Wendelschafer-Crabb, G., Kennedy, W. R., Filla, A., Santoro, L. & Caruso, G. 2001. Small fibers involvement in Friedreich’s ataxia. Annals of Neurology, 50, 17–25.

Piguet, F., De Montigny, C., Vaucamps, N., Reutenauer, L., Eisenmann, A. & Puccio, H. 2018. Rapid and Complete Reversal of Sensory Ataxia by Gene Therapy in a Novel Model of Friedreich Ataxia. Molecular Therapy, 26, 1940–1952.

Puchalska, P. & Crawford, P. A. 2017. Multi-dimensional Roles of Ketone Bodies in Fuel Metabolism, Signaling, and Therapeutics. Cell metabolism, 25, 262–284.

Puchalska, P. & Crawford, P. A. 2021. Metabolic and Signaling Roles of Ketone Bodies in Health and Disease. Annual Review of Nutrition, 41, 49–77.

Ruskin, D. N., Kawamura Jr, M. & Masino, S. A. 2009. Reduced pain and inflammation in juvenile and adult rats fed a ketogenic diet. Plos one, 4, e8349.

Ruskin, D. N., Sturdevant, I. C., Wyss, L. S. & Masino, S. A. 2021. Ketogenic diet effects on inflammatory allodynia and ongoing pain in rodents. Scientific Reports, 11, 1–8.

Sharma, N., Flaherty, K., Lezgiyeva, K., Wagner, D. E., Klein, A. M. & Ginty, D. D. 2020. The emergence of transcriptional identity in somatosensory neurons. Nature, 577, 392–398.

Sourbron, J., Klinkenberg, S., Van Kuijk, S. M., Lagae, L., Lambrechts, D., Braakman, H. M. & Majoie, M. 2020. Ketogenic diet for the treatment of pediatric epilepsy: review and meta-analysis. Child’s Nervous System, 36, 1099–1109.

Szelechowski, M., Amoedo, N., Obre, E., Léger, C., Allard, L., Bonneu, M., Claverol, S., Lacombe, D., Oliet, S., Chevallier, S., Le Masson, G. & Rossignol, R. 2018. Metabolic Reprogramming in Amyotrophic Lateral Sclerosis. Scientific Reports, 8, 3953.

Taylor, M. K., Sullivan, D. K., Mahnken, J. D., Burns, J. M. & Swerdlow, R. H. 2018. Feasibility and efficacy data from a ketogenic diet intervention in Alzheimer’s disease. Alzheimer’s & Dementia: Translational Research & Clinical Interventions, 4, 28–36.

Tierney, J. A., Uong, C. D., Lenert, M. E., Williams, M. & Burton, M. D. 2022. High-fat diet causes mechanical allodynia in the absence of injury or diabetic pathology. Scientific reports, 12, 1–13.

Usoskin, D., Furlan, A., Islam, S., Abdo, H., Lönnerberg, P., Lou, D., Hjerling-Leffler, J., Haeggström, J., Kharchenko, O., Kharchenko, P. V., Linnarsson, S. & Ernfors, P. 2015. Unbiased classification of sensory neuron types by large-scale single-cell Rna sequencing. Nature Neuroscience, 18, 145–153.

Wang, T., Molliver, D. C., Jing, X., Schwartz, E. S., Yang, F.-C., Samad, O. A., Ma, Q. & Davis, B. M. 2011. Phenotypic Switching of Nonpeptidergic Cutaneous Sensory Neurons following Peripheral Nerve Injury. Plos One, 6, e28908.

Wright, D. E., Johnson, M. S., Arnett, M., Smittkamp, S. E. & Ryals, J. M. 2007. Selective changes in nocifensive behavior despite normal cutaneous axon innervation in leptin receptor–null mutant (db/db) mice. Journal of the Peripheral Nervous System, 12, 250–261.

Zhong, S., Zhou, Z., Lin, X., Liu, F., Liu, C., Liu, Z., Deng, W., Zhang, X., Chang, H. & Zhao, C. 2021. Ketogenic diet prevents paclitaxel-induced neuropathic nociception through activation of PPARγ signalling pathway and inhibition of neuroinflammation in rat dorsal root ganglion. European Journal of Neuroscience, 54, 5341–5356.

Zurborg, S., Piszczek, A., Martínez, C., Hublitz, P., Banchaabouchi, M. A., Moreira, P., Perlas, E. & Heppenstall, P. A. 2011. Generation and characterization of an Advillin-Cre driver mouse line. Molecular pain, 7, 1744–8069-7-66.

